# gARID-associated chromatin remodeling events are essential for gametocyte development in *Plasmodium*

**DOI:** 10.1101/2023.06.29.546992

**Authors:** Tsubasa Nishi, Izumi Kaneko, Shiroh Iwanaga, Masao Yuda

## Abstract

Gametocyte development of the *Plasmodium* parasite is a key step for transmission of the parasite from their vertebrate hosts to mosquitoes. Male and female gametocytes are produced from a subpopulation of asexual blood-stage parasites, but the mechanisms that regulate the differentiation of sexual stages are still under investigation. In this study, we investigated the role of gARID, a putative subunit of a chromatin remodeling complex, in transcriptional regulation during the gametocyte development of *P. berghei*. gARID expression starts in early gametocytes before the manifestation of male and female-specific features, and disruption of its gene results in the complete loss of male gametocytes and the production of abnormal female gametocytes. ChIP-seq analysis of gARID showed that it forms a complex with gSNF2, a core subunit of the SWI/SNF chromatin remodeling complex, associating with the male *cis*-regulatory element. Moreover, ChIP-seq of gARID in *gsnf2*-knockout parasites revealed an association of gARID with another *cis*-regulatory element, which is indicated to play a role in both male and female development. Our results showed that gARID functions in two chromatin remodeling events and that remodeling of chromatin states is essential for both male and female gametocyte development.

## Introduction

Malaria is one of the most serious infectious diseases in the world, which is caused by the infection of *Plasmodium* parasites (1). It typically causes fever, vomiting, and headaches, but in severe cases, it can also cause coma, jaundice, seizures, and even death (2). Various strategies, including vector control, chemotherapies, and vaccines, have been used over the last two decades to overcome malaria. However, the disease still occurs in over 200 million individuals and causes 500,000 deaths yearly (3).

*Plasmodium* parasites spread among people through mosquito bites. Thus, the transmission between vertebrate hosts and mosquitoes is an important process for malaria propagation. For the transmission from vertebrates to mosquitoes, the parasites go through sexual development (4, 5). Sexual development begins with a subpopulation of the asexual blood-stage parasites differentiating into male and female gametocytes (6–8). After the gametocytes are ingested by mosquitoes through blood feeding, they egress from red blood cells to form microgamete and macrogamete in the mosquito midgut (9). The gametes then fertilize and develop into ookinetes, which invade the midgut epithelia, reach the basal lamina, and complete the transmission by forming oocysts (10, 11). As the first step of *Plasmodium* sexual development, gametocytogenesis is essential for parasite transmission, and hence, elucidating the mechanism underlying gametocyte development is crucial for developing malaria control strategies such as transmission-blocking drugs and vaccines targeting the sexual stage parasites (12).

*Plasmodium* gametocyte development has been investigated in several studies based on transcriptomics and proteomics (13–19). In addition, recent studies on gametocyte-specific transcription factors significantly contribute to our knowledge of the mechanism underlying gametocyte development, mainly in *P. falciparum* and *P. berghei* (20). Gametocytogenesis is triggered by an AP2-family transcription factor, namely AP2-G (21, 22). It is expressed in a subpopulation of blood-stage parasites, and disruption of its gene results in the complete loss of the ability of parasites to differentiate into gametocytes (23–25). Furthermore, the conversion of parasites into the sexual stage can be induced by the artificial activation of AP2-G (26, 27). After gametocytogenesis is triggered, AP2-G2 induces genome-wide gene repression to support early gametocyte differentiation (28, 29). Moreover, in *P. falciparum*, HDP1 and AP2-G5 were also reported to function during early gametocyte development (30, 31).

The early gametocytes then differentiate into male and female gametocytes. For female gametocyte development, two transcriptional activators, AP2-FG and PFG, regulate the expression of female genes (32, 33). In contrast, two transcriptional regulators, AP2-FG2 and AP2R-2, function together as a transcriptional repressor complex to support female development (34, 35). In addition, transcriptional regulator genes for the zygote and ookinete development, such as *ap2-z* and *ap2-o*, are also transcribed in female gametocytes to prepare for post-fertilization development (36–39).

Compared with that during female gametocyte development, transcriptional regulation during male gametocyte development has been largely unexplored until some recent studies elucidated its mechanisms. A transcriptional regulator, gSNF2, was identified as a core subunit of the SWI/SNF chromatin remodeling complex, which participates in activating male genes by associating with the major male *cis*-regulatory element, TGTCT (40). Hence, the remodeling of the chromatin state is required for male gametocyte development. In addition to *gsnf2*, five male development regulator genes (*md1* through *md5*) were recently identified in *P. berghei* (41). Of these, *md1* was also identified as an essential factor for determining male fate in *P. falciparum* (42). Md1 is a cytoplasmic factor that interacts with ribonucleic granule proteins, and transcription of its gene is considered as a switch to control sex determination.

In our previous study, we reported that in *P. berghei*, one of the roles of AP2-G is to activate transcriptional regulator genes that are important for gametocyte development (43). These genes include most of the abovementioned transcriptional regulator genes. In this study, we investigate one of the transcriptional regulator genes in the AP2-G targets, *garid* (gametocyte-specific AT-rich interactive domain [ARID]-containing protein gene, PBANKA_0102400), during the gametocyte development of *P. berghei*. In a previous study, *garid* was identified as one of the regulator genes essential for male gametocyte differentiation (referred to as *md4*) (41). Moreover, the *P. falciparum* ortholog of gARID was recently identified as a transcriptional regulator essential for microgametogenesis and macrogamete fertility (44). By investigating the role of gARID, we demonstrated that gARID forms a complex with gSNF2 and plays an essential role as a subunit of chromatin remodeling complexes in male differentiation and female development. Furthermore, we revealed that gARID functions in two distinct chromatin remodeling events, and each event is associated with different DNA sequence motifs. We also revealed that the remodeling of the chromatin state is not only important for male gametocyte development but also for female gametocyte development.

## Materials and Methods

### Ethical statement

All experiments in this study were performed according to the recommendations in the Guide for the Care and Use of Laboratory Animals of the National Institutes of Health to minimize animal suffering. All protocols were approved by the Animal Research Ethics Committee of Mie University (permit number 23–29).

### Parasite preparation

All parasites used in this study were inoculated in Balb/c or ddY mice. The *garid*(−), gARID::mNG, and gARID::GFP parasites were derived from the WT ANKA strain, and all the other transgenic parasites were generated via the CRISPR/Cas9 system using Cas9-expressing parasites called PbCas9 (45). The growth rate of blood-stage parasites was assessed by counting infected red blood cells (RBCs) on Giemsa-stained smears every half-day after intraperitoneally injecting infected blood. Ookinete cultures and cross-fertilization assays were performed as previously described (36). Midgut oocysts were counted from the midgut of infected mosquitoes at 14 days post-infective blood meal.

### Generation of transgenic parasites

For tagging gARID with fluorescent proteins, knockout of *garid*, and knockout of *gsnf2* for gARID::GFP^C_*gsnf2*(−)^, the conventional homologous recombination method was used as previously reported (35, 36, 46). Briefly, for tagging experiments, two homologous regions of the *garid* locus were cloned into the *mNG*- or *gfp*-fusion vector to fuse *garid* in-frame with *mNG* or *gfp*. The vector was linearized using restriction enzymes before performing transfection experiments. For knockout experiments, targeting constructs, which contain a *hdhfr* expression cassette flanked with two homologous regions of a gene of interest, were prepared using overlap polymerase chain reaction (PCR). To generate the other transgenic parasites, a previously reported CRISPR/Cas9 system using PbCas9 parasites was used (45). Donor DNAs were constructed using overlap PCR, cloned into pBluescript KS (+), and amplified using PCR. Single guide RNA vectors were prepared by cloning target sequences using annealed oligos.

Transfection experiments were performed using Amaxa Basic Parasite Nucleofector Kit 2 (LONZA). All transfectants were selected by treating mice with 70 μg/mL pyrimethamine, which was added to their drinking water. Recombination was confirmed using PCR for tagging and knockout and using Sanger sequencing for mutations in promoters. Clonal parasites were obtained by limiting dilution. All primers used in this study are listed in Table S5.

### Fluorescence-activated cell sorting (FACS) analysis

FACS analysis was performed using the LSR Fortessa (Becton Dickinson). Nuclei were stained with Hoechst 33342. The analyses were performed using peripheral blood from infected mice with a parasitemia of 2–3%. Cells were gated with forward scatter and Hoechst (450/50) fluorescence intensity. Gated cells were assessed for green fluorescent protein (GFP) (530/30) and red fluorescent protein (RFP) (582/15) fluorescence intensity.

### ChIP-seq and sequencing data analysis

The ChIP-seq experiments were performed as described previously (35). Briefly, parasites were enriched with gametocytes by adding sulfadiazine in the drinking water of infected mice. Whole blood was withdrawn from the infected mice and filtered using Plasmodipur. The blood samples were then fixed with 1% formalin at 30 °C. After fixing, RBCs were lysed in ice-cold 1.5 M NH_4_Cl solution, and the residual cells were lysed in SDS lysis buffer. The lysates were sonicated using Bioruptor (Cosmo Bio) to shear chromatin and subjected to ChIP with antiCGFP polyclonal antibodies (Abcam, ab290) immobilized on Dynabeads Protein A (Invitrogen). DNA fragments purified from the ChIP and input samples were used for library construction using KAPA HyperPrep Kit. The libraries were sequenced using Illumina NextSeq. Two biologically independent experiments were performed for each ChIP experiment.

The obtained sequence data were mapped onto the reference genome sequence of *P. berghei* ANKA, which was downloaded from PlasmoDB 55, using the Bowtie2 tool. Reads aligned at more than two different locations on the genome were removed from the mapping data. From the ChIP and input data, peaks were identified using the macs2 callpeak function with fold enrichment > 3.0 and *q*-value < 0.01, and common peaks between the two experiments were used for further analysis. Binding motifs were predicted by analyzing the enrichment of motifs within 50 bp from peak summits using Fisher’s exact test. Genes that possessed peaks within the 1,200-bp upstream region from ATG were identified as target genes. Parameters for all programs were set to the default unless specified otherwise.

### RNA-seq and sequence data analysis

RNA extraction and RNA-seq experiments were performed as described previously (35). Briefly, gametocytes were enriched in the host as described above, and total RNA was extracted from the Plasmodipur-filtered whole blood of the infected mice using the Isogen II reagent (Nippon gene). From the total RNA, RNA-seq libraries were prepared using the KAPA mRNA HyperPrep Kit and sequenced using Illumina NextSeq. Three biologically independent experiments were performed for each sample. The obtained sequence data were mapped onto the reference genome sequence of *P. berghei* by HISAT2, setting the maximum intron length threshold to 1,000. The mapping data for each sample were analyzed using featureCounts and compared using DESeq2. The fragments per kilobase of transcript per million mapped reads (FPKM) for each gene were calculated from the featureCounts data, and genes with FPKM < 10 in all three datasets for WT were removed before the differential expression analysis. Genes in subtelomeric regions were also removed. For all programs, the parameters were set to the default unless specified otherwise.

## Result

### gARID is a putative subunit of a chromatin remodeling complex conserved in Apicomplexa

*garid* (PBANKA_0102400) is a target gene of AP2-G (43), and its transcription is upregulated via conditional induction of AP2-G in *P. falciparum* and *P. berghei* (26, 27). *garid* encodes a protein that contains an ARID at its N-terminus and a monopartite nuclear localization signal near its C-terminus (Fig 1A). The ARID is a helix–turn–helix motif-based DNA binding domain conserved in a wide variety of eukaryotic species from fungi to vertebrates (47, 48). In mammals, proteins with this domain are classified into 7 subfamilies, namely ARID1 through ARID5, JARID1, and JARID2 (49). Of these, only ARID3 and ARID5 preferentially interact with AT-rich sequences, and the others have no clear sequence preference (50). These ARID proteins participate in the regulation of cell growth, differentiation, and development. To investigate which ARID subfamily gARID can be classified into, we performed a protein–protein BLAST (blastp) search within “homo sapiens” using the ARID domain (95 residues from the N-terminus) of gARID as a query. The blastp result showed that the ARID domain of gARID matched best with human ARID2 with an E-value of 7.0 × 10^−6^ (Fig 1B). Next, we constructed a dendrogram of ARID domains from gARID and ARID family proteins in humans and *Drosophila*, the two species in which ARID proteins have been extensively characterized. The tree showed that gARID is grouped into a clade with ARID2 and BAP170 (an orthologue of ARID2 in Drosophila), separate from the other ARID proteins (Fig 1C). ARID2 is a subunit of the chromatin remodeling complex, polycomb-associated BRG1/BRM-associated factor (PBAF), which is a type of SWI/SNF complex (51, 52). These results suggested that gARID could be related to the ARID2 subfamily.

**Fig 1.**
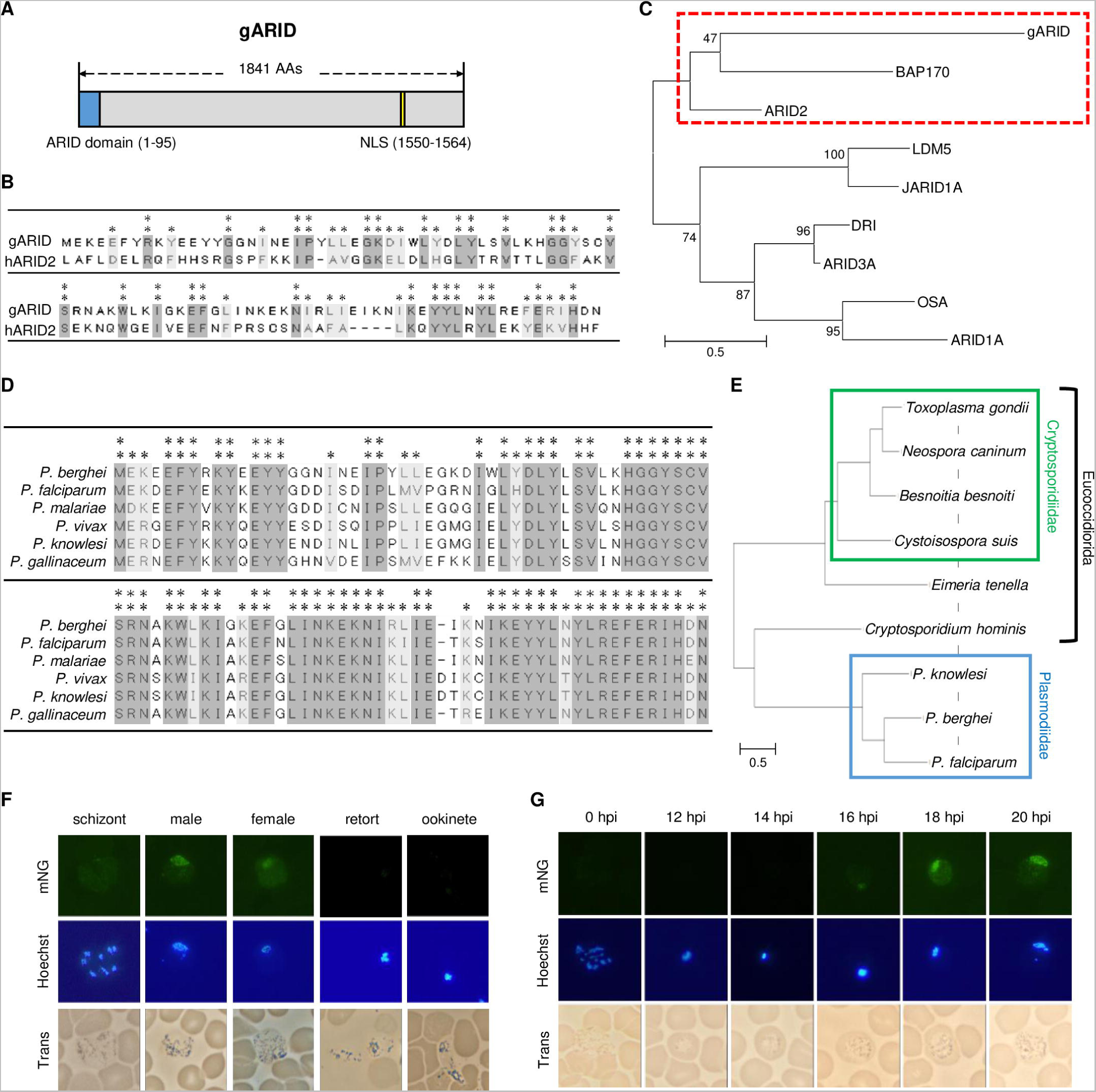
Features of gARID and its expression during gametocyte development. (A) A schematic illustration of gARID. The blue box shows the AT-rich interactive domain (ARID) domain. The nuclear localization signal (yellow box) was predicted using cNLS Mapper (http://nls-mapper.iab.keio.ac.jp/cgi-bin/NLS_Mapper_form.cgi). Amino acid positions for each feature are shown in brackets. (B) Alignment of amino acid sequences of gARID and human ARID2 (hARID2). ** represents positions at which both sequences possess the same amino acid, whereas * represents positions with amino acid residues with similar properties. (C) A dendrogram of ARID domains from gARID and ARID family proteins in human and *Drosophila* (human ARID [ARID1A, ARID2, ARID3A, and JARID1A] and *Drosophila* ARID [OSA, BAP170, DRI, and LDM5]). Each ARID domain was determined using the simple modular architecture research tool. The amino acid sequences of ARID domains were aligned using the ClustalW program in Mega X. The tree was inferred from the alignment using the maximum likelihood method and Jones–Taylor–Thornton matrix-based model and was drawn to scale, with branch lengths measured according to the number of substitutions per site. The numbers above branches show the bootstrap values (1,000 replicates). (D) Alignment of amino acid sequences of ARID domain from gARID orthologs in *Plasmodium* species by the ClustalW program in Mega X (*Plasmodium berghei*, PBANKA_0102400; *Plasmodium falciparum*, PF3D7_0603600; *Plasmodium malariae*, PmUG01_11059200; *Plasmodium vivax*, PVP01_1145300; *Plasmodium knowlesi*, PKNH_1147100; and *Plasmodium gallinaceum*, PGAL8A_00132100). (E) Phylogenetic tree of gARID and its putative orthologs in Apicomplexan parasites (*P*. *berghei*, gARID; *P*. *falciparum*, PF3D7_0603600; *P*. *knowlesi*, PKNH_1147100; *Cryptosporidium hominis*, XP_665701; *Eimeria tenella*, XP_013235762; *Toxoplasma gondii*, XP_018634911; *Neospora caninum*, XP_003885918; *Cystoisospora suis*, PHJ24466; and *Besnoitia besnoiti*, XP_029222326). The whole amino acid sequence for each protein was used for alignment. The tree was inferred as described in (C). (F) gARID expression in asexual and sexual stages of the gARID::mNG. Asexual-blood stages and gametocytes were assessed on peripheral blood smears. Retort-form and banana-shaped ookinetes were assessed in ookinete cultures at 16 and 20 h after starting the cultures, respectively. Nuclei were stained with Hoechst 33342. (G) Time-course fluorescent analysis of the gARID::mNG during gametocyte development. Synchronized schizonts before injection were described as parasites at 0 h post-injection (hpi).

The ARID domain of gARID is highly conserved in *Plasmodium* species (Fig 1D). To evaluate whether gARID is conserved in other Apicomplexan parasites, we performed the blastp search using the ARID domain of gARID. The search identified proteins with an ARID domain from apicomplexan parasites such as *Toxoplasma* and *Cryptosporidium* (Fig S1). Notably, the amino acids conserved among gARID and the blastp-detected proteins contained most of the consensus amino acids for ARID domains. The phylogenetic tree of gARID and the blastp-detected proteins was topologically consistent with the species tree of Apicomplexa, except that *Cryptosporidium* was supposed to be closer to the clade of Sarcocystidae and Eimeriidae than that of Plasmodiidae (Fig 1E). This result further suggested that these blastp-detected proteins are an ortholog of gARID.

### gARID is specifically expressed from early to mature gametocyte development

To investigate the cell stages in which gARID functions, we generated a parasite line that expresses gARID fused with mNeon Green (gARID::mNG, Fig S2A). No fluorescent signal was detected in any asexual stage parasites when the fluorescent analysis was performed using gARID::mNG (Fig 1F). In contrast, nuclear-localized signals were detected in both male and female gametocytes (Fig 1F). We further assessed the fluorescence in ookinete cultures and observed no fluorescent signal in ookinetes (Fig 1F).

To evaluate the expression pattern of gARID in detail during gametocyte development, we performed fluorescent analyses in a time-course manner using cell cycle-synchronized parasites. We intravenously injected mice with mature schizonts, which were cultured *in vitro* and collected via density gradient centrifugation, and observed fluorescence of gARID::mNG every 2 h. Parasites showed no fluorescent signal from 0 h post-injection (hpi), *i.e.*, schizonts before injection, to 14 hpi (Fig 1G). At 16 hpi, at which no parasites have yet exhibited any sex-specific features, a small population of parasites started to express nuclear-localized fluorescence, and the signal became stronger at 18 hpi (Fig 1G). This result indicated that the expression of gARID starts in early gametocytes before differentiation into male or female gametocytes. The fluorescent signals then continued until the maturation of gametocytes. This expression pattern of gARID is consistent with the fact that *garid* is a target gene of AP2-G, because our previous result showed that the expression of AP2-G starts at 12–14 hpi in *P. berghei*.

### Disruption of *garid* results in the complete loss of male gametocyte production

To investigate the role of gARID in gametocyte development, we disrupted *garid* through a conventional homologous recombination method using a pyrimethamine resistance marker (*garid*[−], Fig S2B). Two clonal lines were obtained from independent transfection experiments and used for phenotype analyses. The *garid*(−) parasites exhibited a growth rate that was comparable to that of the parental *Pb*ANKA strain (WT) (Fig 2A), indicating that gARID plays no role in asexual blood-stage development. We next assessed the production of gametocytes by observing parasites in Giemsa-stained blood smears. In the assay, female gametocytes were observed, but no male gametocytes were observed in *garid*(−) (Fig 2B). Consistently, no exflagellation was detected in *garid*(−) after the activation of gametogenesis by exposing parasites to an ookinete culture medium (pH 8.2 and 20 °C). As no male gametocyte was observed in the peripheral blood, *garid*(−) could not form any zygotes when cultured in an ookinete culture medium for both clones (Fig 2C). In addition, no oocysts were observed in the mosquito midgut 14 days after the blood meal (Fig 2D). To further confirm the absence of male gametocytes in *garid*(−), we knocked out *garid* in the gametocyte reporter line 820cl1 (*garid*[−]^820^), which have GFP and RFP expression cassettes under the control of male- and female-specific promoters, respectively. FACS analysis detected no GFP-positive parasites in the *garid*(−)^820^ parasites compared with the parental 820cl1 wherein approximately 3% of the parasites showed a GFP signal (Fig 2E). This confirmed that disruption of *garid* results in the complete loss of male production.

**Fig 2.**
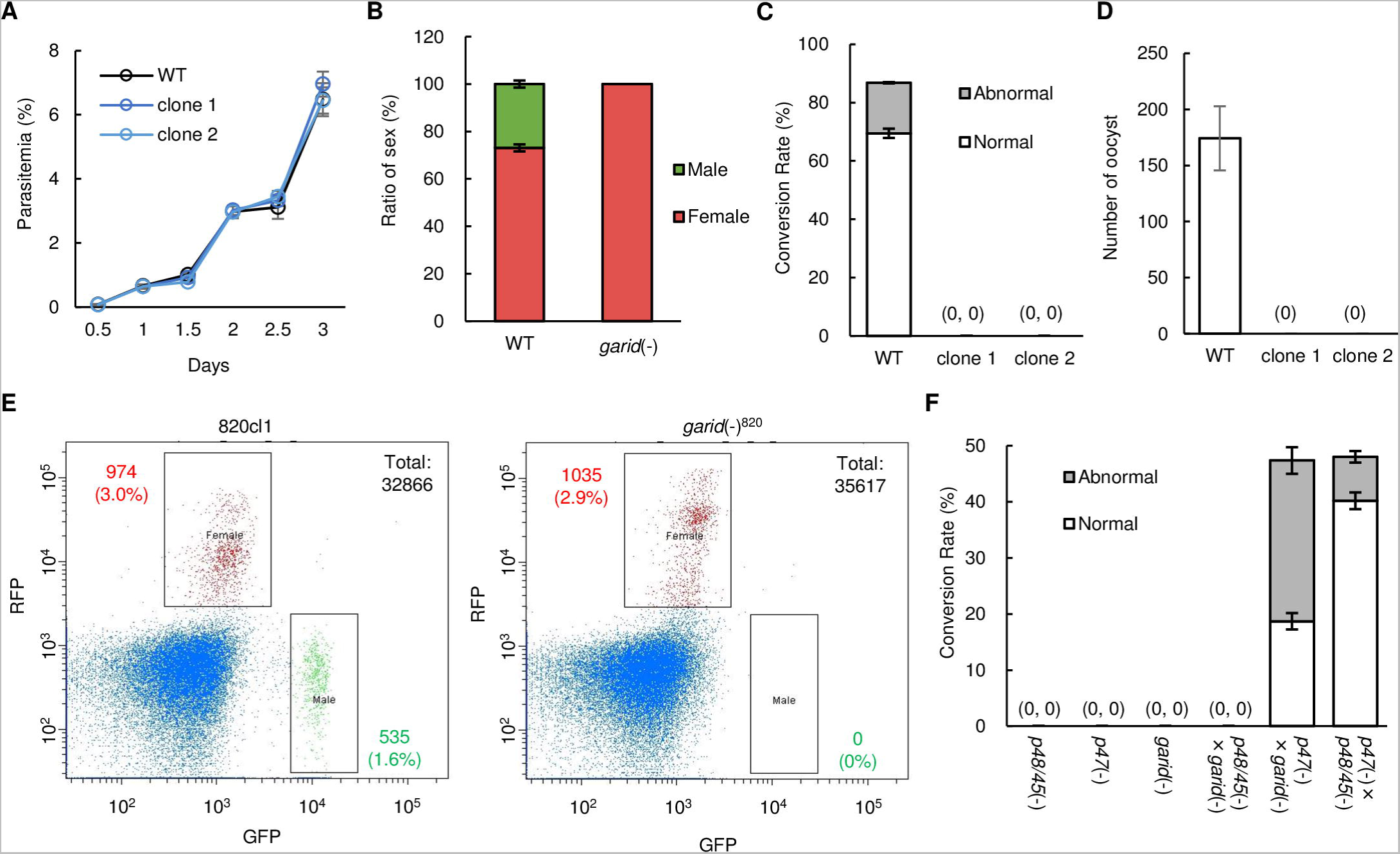
Knockout phenotype of *garid*. (A) Growth rate of blood-stage in wild type (WT) and *garid*(−) parasites. Parasitemia was calculated by counting infected red blood cells on Giemsa-stained blood smears. The date for injection of infected blood into mice was set as day 0. (B) The ratio of male and female gametocytes in WT and *garid*(−) parasites. The number of gametocytes was assessed on Giemsa-stained blood smears in the gametocyte-enriched condition. Error bars indicate the standard error of the mean (n = 3). (C) The conversion rate of female gametocytes to normal ookinetes (banana-shaped; white box) and abnormal ookinetes (retort-form ookinetes and fertilized cells without apical protrusion; grey box). The number of female-derived cells was counted at 20 h after starting ookinete cultures. Error bars indicate the standard error of the mean (n = 3). (D) The number of midgut oocysts in WT and *garid*(−) parasites at 14 days post-infection. Error bars indicate the standard error of the mean (n = 20). (E) Fluorescence-activated cell sorting analysis of 820cl1 and *garid*(−)^820^ showing the numbers of female and male gametocytes as red and green fluorescent protein-positive parasites, respectively. Cells gated with forward-scatter and Hoechst are shown in the plot, and the number is indicated as “Total” in the top right corner. Nuclei were stained with Hoechst 33342. (F) Cross-fertilization assay using *garid*(−), *p48/45*(−), and *p47*(−). The *p48/45*(−) and *p47*(−) parasites are male- and female-defective, respectively. The conversion rate for normal and abnormal ookinetes are shown in white and grey boxes, respectively. Error bars indicate the standard error of the mean (n = 3).

We then performed cross-fertilization assays to evaluate whether female gametocytes of *garid*(−) can develop into ookinete when fertilized with normal male gametocytes. When *garid*(−) was crossed with parasites that produce infertile males (*p48/45*[−]) (53), no zygotes were observed, which was consistent with the absence of male gametocytes in *garid*(−) (Fig 2F). In contrast, when *garid*(−) was crossed with parasites that produce infertile females (*p47*[−]) (54), parasites could undergo fertilization at a ratio similar to that when *p48/45*(−) and *p47*(−) were crossed (Fig 2F). However, the ratio of conversion from female to banana-shaped ookinetes was only 19%, which was significantly smaller than the ratio for crossing *p48/45*(−) and *p47*(−), which was 40% (*p*-values = 4.9 × 10^−4^ using two-tailed Student’s *t*-test) (Fig 2F). This result suggested that although female gametocytes of *garid*(−) showed normal morphology, they exhibit an abnormality that causes delay or arrest of ookinete development after fertilization. Collectively, these results indicated that gARID plays a role in differentiation into male gametocytes and the development of female gametocytes. These results for the phenotype analyses of *garid*(−) are mostly consistent with a recent study by Russell *et al.*, except that the female gametocytes of the *garid*-knockout parasite did not show any defects based on their data (41). Meanwhile, in *P. falciparum*, disruption of the *garid* ortholog affects female gamete fertility, consistent with our result (44).

### gARID is essential for regulating gametocyte transcriptome

Because gARID is nuclear-localized, we hypothesized that gARID is involved in the transcriptional regulation of gametocytes. Accordingly, we performed a differential expression analysis between the gametocytes of WT and *garid*(−) to investigate its role in transcriptional regulation. Total RNA was harvested from gametocyte-enriched parasites by killing asexual blood-stage parasites using sulfadiazine treatment and used for RNA-seq analysis. Three independent samples were analyzed for each strain, and the sequence data were compared using DESeq2 (55). The analysis showed that in *garid*(−), 264 genes were significantly downregulated (log_2_[fold change] < −1, *p*-value adjusted for multiple testing with the Benjamini-Hochberg procedure [*p*-value adj] < 0.05), whereas only 15 genes were significantly upregulated (log_2_[fold change] > 1, *p*-value adj < 0.05) compared with those in WT (Fig 3A and Table S1).

**Fig 3.**
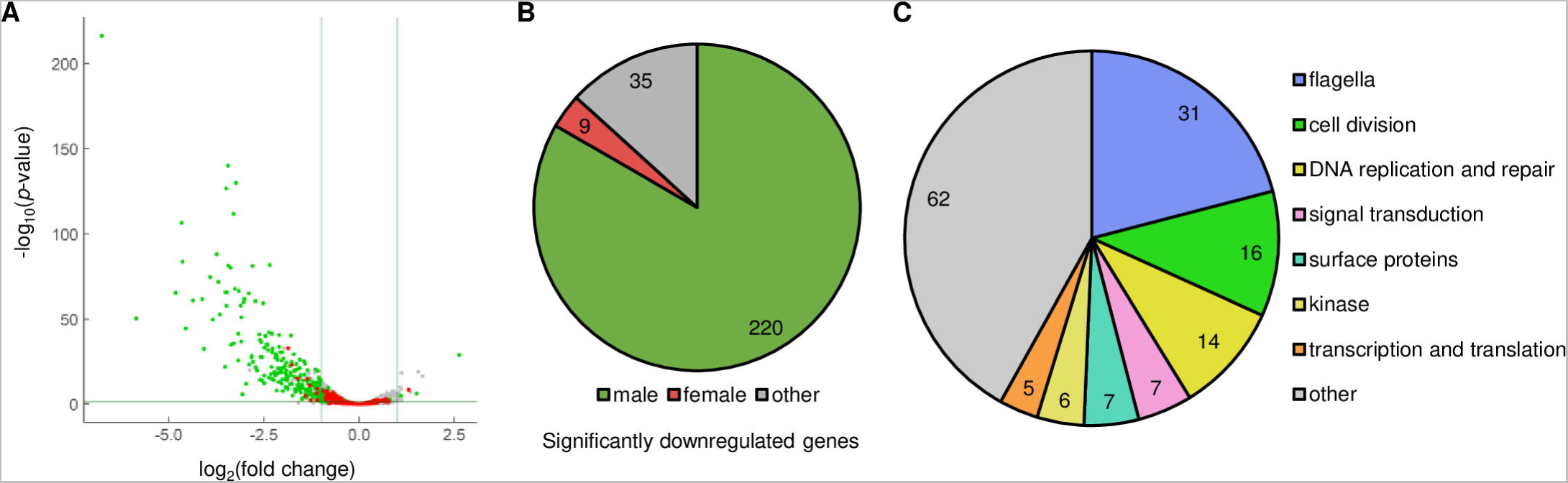
Differential expression analysis between wild-type (WT) and *garid*(−) gametocytes. (A) Volcano plot showing differential gene expression between WT and *garid*(−) gametocytes. Green and red dots represent male- and female-enriched genes, respectively. The horizontal line indicates a *p*-value of 0.05, and the two vertical lines indicate a log_2_(fold change) of 1 and −1. (B) Classification of genes, which were significantly downregulated in *garid*(−), into male- and female-enriched genes and others. (C) Classification of the significantly downregulated genes that are functionally annotated on PlasmoDB into seven functional groups and “other.”

We then evaluated the effects of disrupting *garid* on male and female gametocyte transcriptome by comparing the significantly downregulated genes with the sex-enriched genes, which were previously identified using the sex-specific transcriptome data (19, 35). Of the 264 significantly downregulated genes, 220 genes were male-enriched, which was consistent with the absence of male gametocytes in *garid*(−) (Fig 3B). They accounted for more than half of male-enriched genes (220/438) and included well-known male genes, such as *hap2*, *MiGS*, *p230,* and *p230p* (54, 56–59). In addition, many of the downregulated genes could be classified into the functional groups of major male features; in the classification of 148 downregulated genes that are functionally annotated on PlasmoDB (https://plasmodb.org/plasmo/app), the three largest groups were “flagella,” “DNA replication and repair,” and “cell division” (Fig 3C). Along with the phenotype analysis, these results suggested that gARID functions as a major factor for male gene activation.

### gARID directly activates a majority of male genes

To investigate whether gARID directly regulates the genes downregulated in *garid*(−), we performed chromatin immunoprecipitation followed by high-throughput sequencing (ChIP-seq) analysis using parasites that express GFP-fused gARID (gARID::GFP, Fig S2C) and anti-GFP antibody. The IPed DNA fragments were sequenced via next-generation sequencing (NGS), and peaks were identified from the sequence data. Two independent experiments were performed, and 1263 and 1258 peaks were identified in Experiments 1 and 2, respectively (Table S2A and S2B). Of these peaks, 1115 peaks overlapped between the two experiments, indicating high reproducibility of the ChIP-seq experiments (Fig 4A).

**Fig 4.**
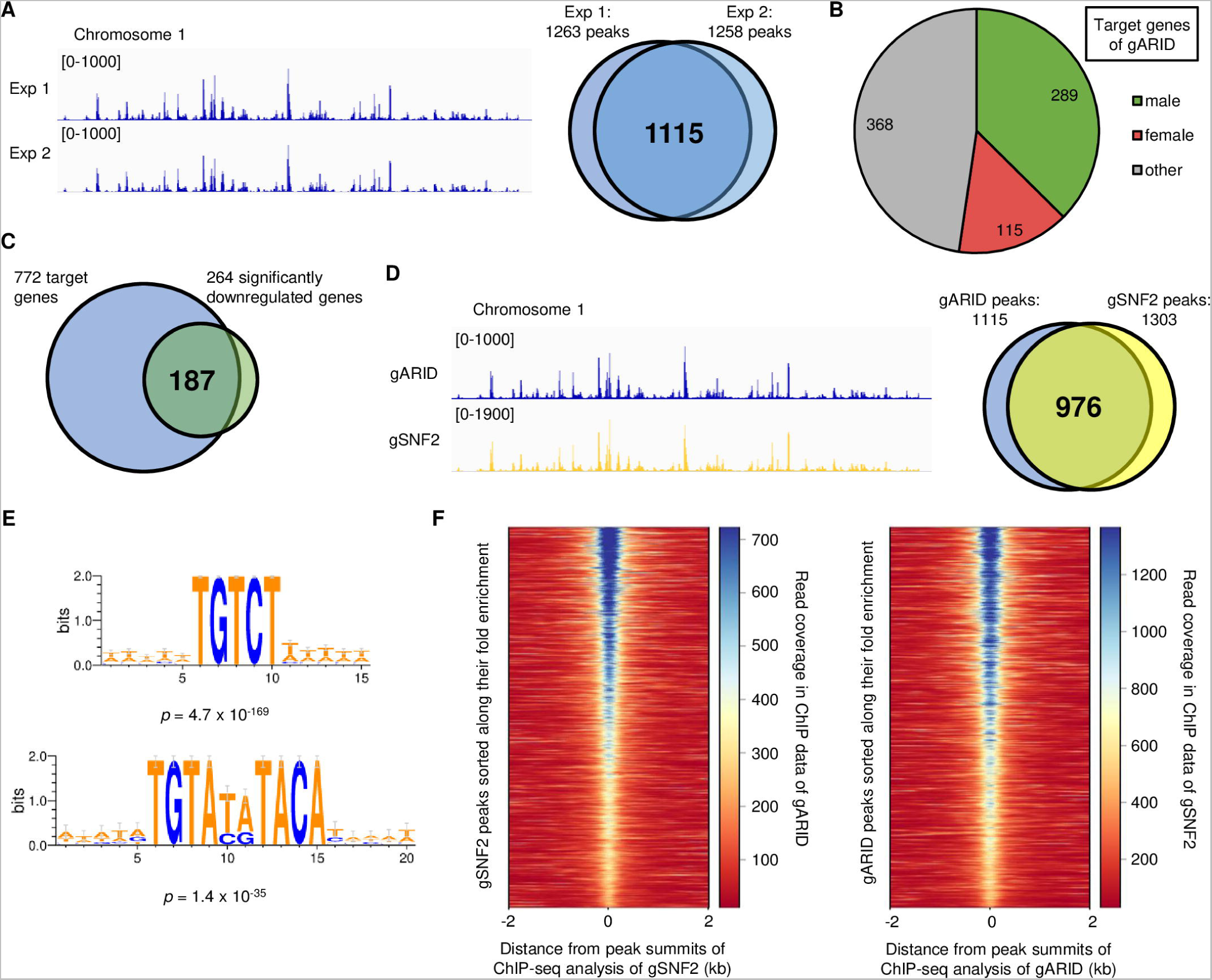
ChIP-seq analysis of gARID. (A) Integrative Genomics Viewer (IGV) images from ChIP-seq Experiments 1 and 2 of gARID on the whole chromosome 1. Histograms show row read coverage of the ChIP data at each base. The scales are shown in brackets. The Venn diagram on the right shows the number of peaks overlapped between Experiments 1 and 2. (B) Classification of gARID target genes into sex-enriched gene sets. (C) Venn diagram showing overlap of the gARID target genes and the significantly downregulated genes in *garid*(−). (D) IGV images showing ChIP-seq peaks of gARID and gSNF2 on the whole of chromosome 1. The scales are shown in brackets. The Venn diagram on the right shows an overlap of peaks identified in ChIP-seq of gARID and gSNF2. (E) Enrichment of TRTAYRTACA and TGTCT motifs within 50 bp from peak summits identified in the ChIP-seq of gARID. The logos were depicted by WebLogo 3 (http://weblogo.threeplusone.com/). (F) Heatmaps showing coverage in ChIP-seq of gARID at gSNF2 peaks (left) and that of gSNF2 at gARID peaks (right). Peaks are aligned in ascending order of their fold enrichment values from the top of heat maps.

From the Chip-seq data, we searched for genes with a peak summit located within 1.2 kbp upstream from the start codon and defined them as a target gene of gARID. Through the search, we identified 772 target genes (Table S2C). These target genes comprised the majority of male-enriched genes (289/444 male-enriched genes), indicating the role of gARID in directly regulating male genes (Fig 4B). The targets also contained a considerable number of female-enriched genes, suggesting that gARID plays a role in female development as well, which is consistent with the results of the phenotype analysis (Fig 4B). We then investigated whether these target genes were differentially expressed in *garid*(−) and revealed that the targets were enriched in the significantly downregulated genes (187/264 genes) with *p*-values of 2.5 × 10^−85^ using Fisher’s exact test (Fig 4C). This result indicated that gARID is involved in the transcriptional activation of its target genes.

Notably, the genome-wide peak pattern of the ChIP-seq of gARID was almost identical to that of gSNF2, which we had previously reported (Fig 4D) (40). gSNF2 is a core subunit of SWI/SNF2 chromatin remodeling complex expressed in gametocytes, and *gsnf2* knockout parasite (*gsnf2*[−]) showed impaired male gametocyte development (40). Of the peaks identified in the ChIP-seq of gARID and gSNF2, 976 peaks overlapped between them, which accounted for nearly 90% of the ChIP peaks of gARID (Fig 4D). Consistently, we found the enrichment of the two motifs, TGTCT and TGTAYRTACA, which were gSNF2-associated motifs, within 50 bp of the peak summits in the gARID ChIP-seq with *p*-values of 4.7 × 10^−169^ and 1.4 × 10^−35^ using Fisher’s exact test, respectively (Fig 4E). Moreover, the intensity of each corresponding peak was highly consistent between the two ChIP-seq data; when a heatmap of read coverage for the ChIP data of gARID was depicted, the centering location of the peak summits identified in the ChIP-seq of gSNF2, higher read counts in gARID ChIP-seq was observed at the gSNF2 peaks with higher fold enrichment (Fig 4F, left). The same was true for the ChIP-seq coverage of gSNF2 at the gARID peaks (Fig 4F, right). Together with the results in the protein sequence analysis of ARID domains that indicated the relationship between gARID and ARID2, it was strongly suggested that gARID forms a complex with gSNF2 on the genome and together functions as the SWI/SNF2 chromatin remodeling complex.

### gARID plays an additional role during gametocyte development independently from gSNF2

The ChIP-seq analysis of gARID indicated the cooperation of gARID and gSNF2 in the transcriptional regulation of their target genes. However, their knockout phenotypes are not the same; *gsnf2*(−) produces immature male gametocytes, whereas *garid*(−) completely loses the ability to differentiate into male gametocytes. In addition, female gametocyte development was affected in *garid*(−) but not in *gsnf2*(−). Consistent with the differences in their phenotype, the results in the differential expression analyses for *garid*(−) vs. WT and *gsnf2*(−) vs. WT were also different; the majority of male- and female-enriched genes were more downregulated in *garid*(−) than in *gsnf2*(−) (Fig 5A and 5B). Furthermore, a direct comparison of the RNA-seq data between *garid*(−) and *gsnf2*(−) identified 516 genes that were significantly downregulated in *garid*(−) compared with those in *gsnf2*(−), which contained 94 male- and 112 female-enriched genes (Table S3). However, compared with those in WT, the downregulation of female-enriched genes was barely detected in *garid*(−). This was probably because the significant difference in the male transcriptome between WT and *garid*(−) masked the less significant changes in the female transcriptome. Compared with those in *gsnf2*(−), the downregulation of female-enriched genes in *garid*(−) only became detectable due to less significant differences in their male transcriptomes.

**Fig 5.**
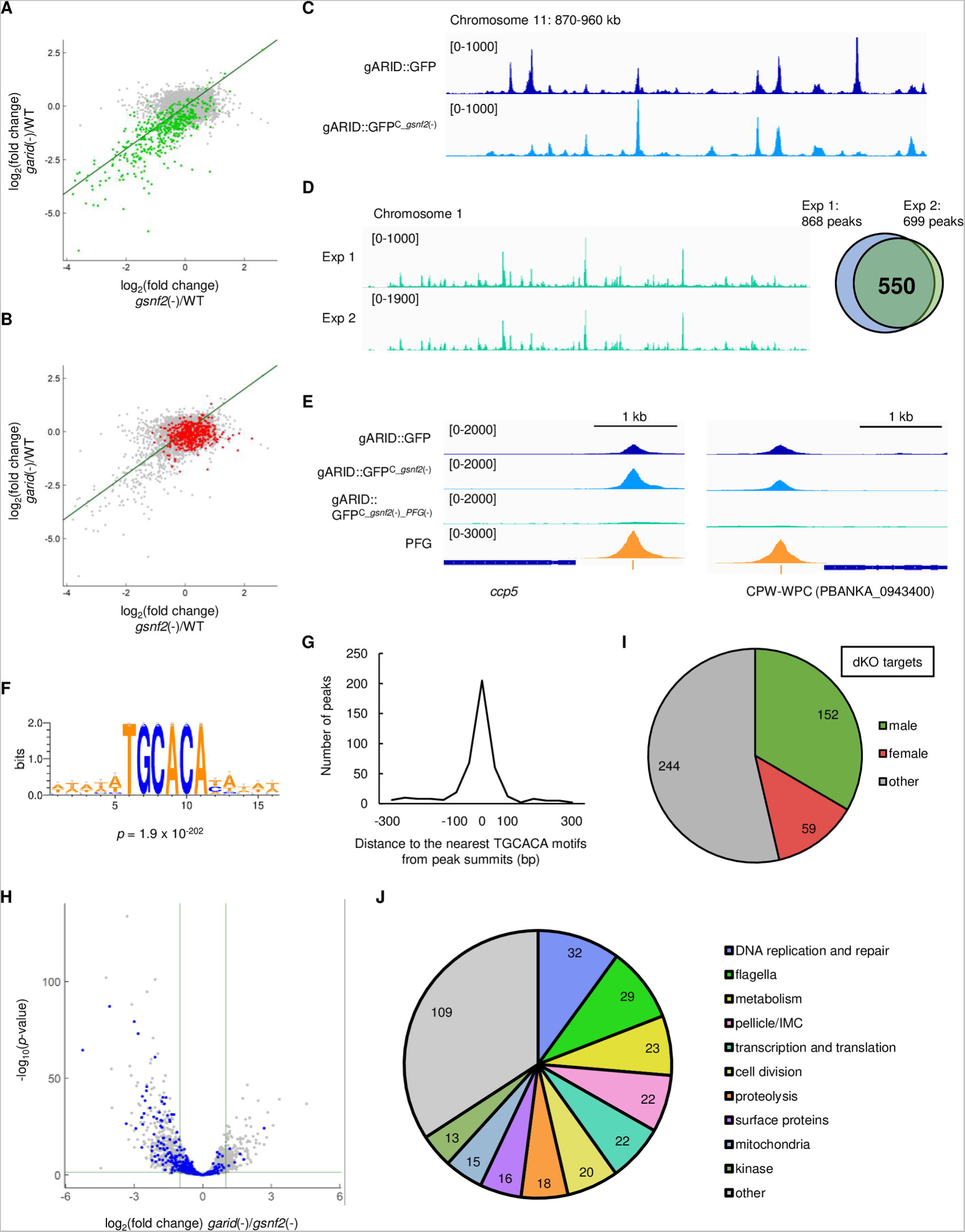
Function of gARID in the absence of gSNF2. (A) The scatter plot showing the relationship between log_2_(fold change) for *garid*(−) vs. wild type (WT) and *gsnf2*(−) vs. WT for each gene. Green dots represent male-enriched genes. The line shows a slope of 1 and an intercept of 0. (B) The scatter plot showing data mentioned in (A) for female-enriched genes, as indicated by red dots. (C) Integrative Genomics Viewer (IGV) images showing ChIP-seq peaks of gARID in gARID::GFP and gARID::GFP^C_*gsnf2*(−)^ on a part of chromosome 11. Histograms show row read coverage of ChIP data at each base. The scales are shown in brackets. (D) IGV images from ChIP-seq Experiment 1 and 2 using gARID::GFP^C_*gsnf2*(−)_*pfg*(−)^ on the whole chromosome 1. The scales are shown in brackets. The Venn diagram on the right shows the number of peaks overlapped between Experiments 1 and 2. (E) IGV images showing peaks that disappeared after *pfg* disruption. ChIP-seq peaks of gARID in gARID::GFP, gARID::GFP^C_*gsnf2*(−)^, and gARID::GFP^C_*gsnf2*(−)_*pfg*(−)^ and peaks of PFG are shown. Yellow bars indicate the positions of TGTAYRTACA motifs. The scales are shown in brackets. (F) A DNA motif enriched within 50 bp from peak summits identified in the ChIP-seq using gARID::GFP^C_*gsnf2*(−)_*pfg*(−)^. The logo was depicted by WebLogo 3. (G) Distance between peak summits identified in the ChIP-seq using gARID::GFP^C_*gsnf2*(−)_*pfg*(−)^ and the nearest TGCACA motifs. (H) Volcano plot showing differential gene expression between gametocytes of *garid*(−) and *gsnf2*(−). Blue dots represent the gARID target genes in gARID::GFP^C_*gsnf2*(−)_*pfg*(−)^. The horizontal line indicates a *p*-value of 0.05, and the two vertical lines indicate log_2_(fold change) of 1 and −1. (I) Classification of gARID target genes in gARID::GFP^C_*gsnf2*(−)_*pfg*(−)^ into sex-enriched gene sets. (G) Classification of functionally annotated target genes of gARID in gARID::GFP^C_*gsnf2*(−)_*pfg*(−)^ into ten functional groups and “other.”

From this result, we considered that gARID plays an additional role in transcriptional regulation during gametocyte development. To explore the interaction of gARID with factors other than gSNF2, we performed a ChIP-seq of gARID in the *gsnf2*-knockout parasite. For this purpose, gARID::GFP was generated with the CRISPR/Cas9 system using Pbcas9 (gARID::GFP^C^, Fig S3A), and then, *gsnf2* was disrupted in the gARID::GFP^C^ (gARID::GFP^C_*gsnf2*(−)^, Fig S3B). The ChIP-seq analysis showed a different pattern of peaks compared with that in the ChIP-seq using gARID::GFP (Fig 5C). Around these peak summits, the TGTCT motif was no longer enriched (*p*-value = 0.25). This result confirmed that gARID is recruited by gSNF2 as a subunit of the SWI/SNF complex on the TGTCT motifs. In contrast, the enrichment of the TGTAYRTACA was still observed with a *p*-value of 8.7 × 10^−104^ using Fisher’s exact test. In our previous study, we showed that the female-specific transcriptional activator, PFG, is responsible for recognizing the TGTAYRTACA motifs (33). Therefore, gARID is recruited to the TGTAYRTACA motifs by PFG independent of gSNF2. In addition to the ten-base motif, the TGCACA motif was also enriched around the peak summits identified in the ChIP-seq using gARID::GFP^C_*gsnf2*(−)^ (*p*-value = 3.0 × 10^−117^), suggesting that it could be another important gARID-associated sequence motif.

To thoroughly investigate the property of gARID binding regions other than those on the known motifs, we knocked out both *gsnf2* and *pfg* in gARID::GFP^C^ using CRISPR/Cas9 (gARID::GFP^C_*gsnf2*(−)_*pfg*(−)^, Fig S3C and S3D) and performed ChIP-seq analysis. Two independent experiments identified 868 and 699 peaks, of which 550 peaks overlapped between the two experiments (Fig 5D, Table S4A, and S4B). As expected, peaks around the TGTAYRTACA motif mostly became undetectable after the disruption of *pfg*, and its enrichment in peak regions was significantly decreased (*p*-value = 7.5 × 10^−3^) (Fig 5E). However, the TGCACA motif was significantly enriched in the peak regions with a *p*-value of 1.9 × 10^−202^ using Fisher’s exact test (Fig 5F). In addition, the TGCACA motif was found within 300 bp from the summit of 424 peaks (77% of the peaks), and most of them were located within 100 bp from the summit (Fig 5G). These results suggested that gARID is indeed associated with the TGCACA motif. We further assessed genome-wide peak location and identified 455 target genes of gARID in gARID::GFP^C_*gsnf2*(−)_*pfg*(−)^ (dKO targets, Table S4C). The dKO targets included 102 genes that were significantly downregulated in *garid*(−) compared with those in *gsnf2*(−), with a *p*-value of 2.0 × 10^−11^ using Fisher’s exact test (Fig 5H). In contrast, the genes downregulated in *gsnf2*(−) compared with those in *garid*(−) only contained seven of the dKO targets. Therefore, we considered that as the additional role, gARID activates genes downstream of the TGCACA motif, from which the phenotypic differences in the gametocyte development between *garid*(−) and *gsnf2*(−) could be derived.

To evaluate the role of gARID on the TGCACA motif in gametocyte development, we classified the 319 functionally annotated dKO targets into 10 functional groups and “other.” The groups for male-related functions, “DNA replication and repair” and “flagella,” were the two major groups that contained 32 and 29 genes, respectively (Fig 5J). Furthermore, another male-related functional group, “cell division,” also contained a significant number of genes. Most of these male genes are also included in the targets of the TGTCT motifs. Therefore, this result suggested that some male genes are regulated by the following two different complexes containing gARID: one with gSNF2 on the TGTCT motif and the other on the TGCACA motif, probably with an unknown remodeler. In addition to these male-related genes, the dKO targets also contained several genes in female-related groups, such as “pellicle/IMC” and “mitochondria.” Moreover, the targets included a considerable number of both sex-enriched genes, *i.e.*, 152 male- and 59 female-enriched genes (Fig 5I), implying the role of the TGCACA motif in both male and female gametocyte development.

### The TGCACA motif functions as a *cis*-regulatory element for activating male genes

To investigate the function of the TGCACA motif as a *cis*-regulatory element and the relationship between the TGTCT and TGCACA motifs in male gametocyte development, we performed reporter assays using endogenous loci. As a target for the reporter assays, we selected the following two genes: *calm* (PBANKA_1421000, a putative calmodulin gene) as a gene significantly downregulated in *garid*(−) but only slightly in *gsnf2*(−) (log_2_[fold change] = −2.3 and −0.92, respectively) and *rsph9* (PBANKA_1431500, a gene encoding a putative radial spoke head protein 9 homolog, a component of radial spokes which control axonemal dynein activity) (60–62) as a gene downregulated in both *garid*(−) and *gsnf2*(−) (log_2_[fold change] = −6.8 and −3.6, respectively) (Table). We first tagged each of these genes with *gfp* to assess their expression using FACS (CALM::GFP and RSPH9::GFP, Fig S4A and S4B). Then, we mutated the TGTCT and TGCACA motifs, either or both, within the peak regions located upstream of *calm* and *rsph9* to assess the roles of these two *cis*-regulatory elements in the activation of male genes (Fig 6A, S4C, and S4D).

**Fig 6.**
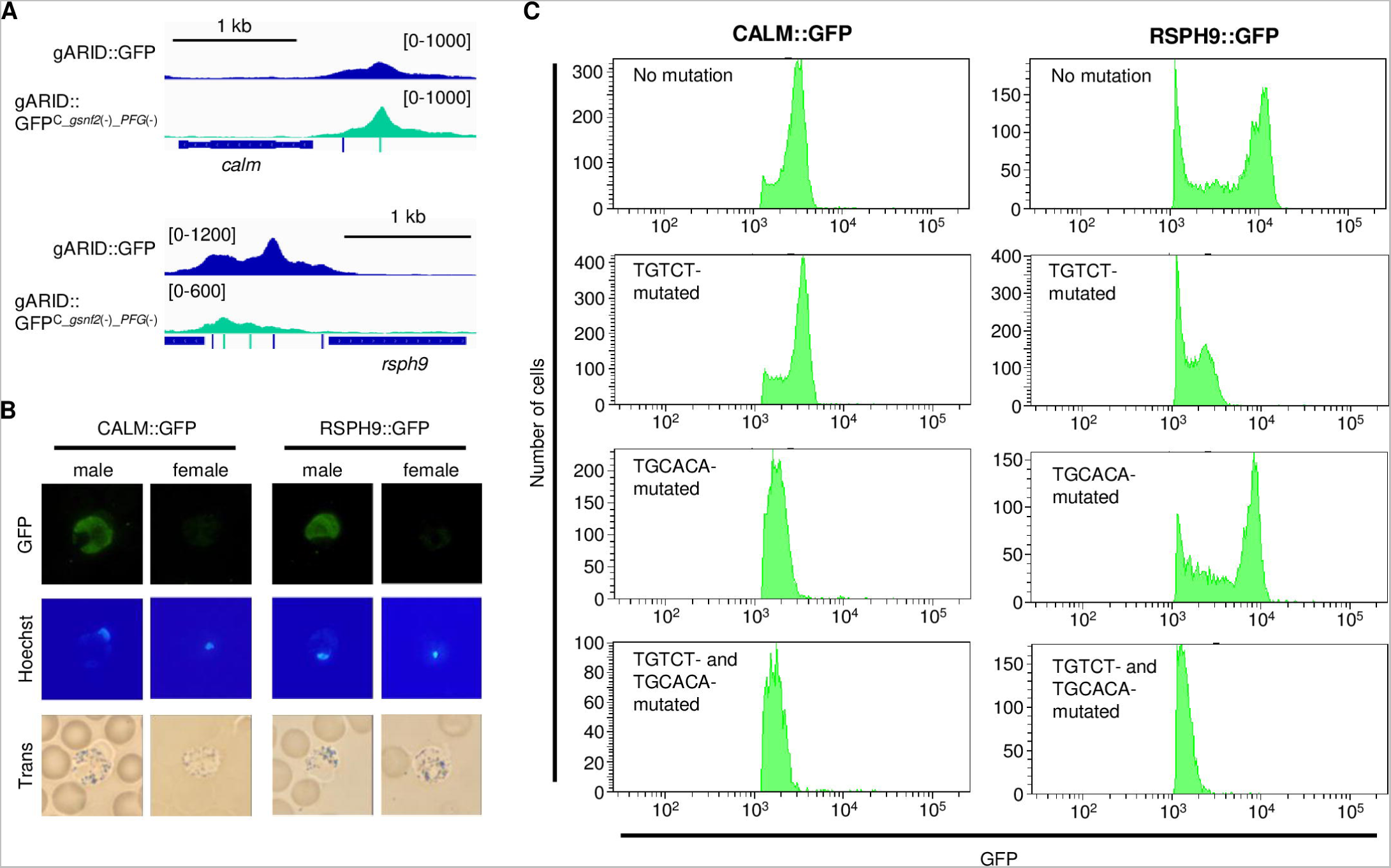
Role of the TGCACA motif as a *cis*-regulatory element for activating male genes. (A) ChIP-seq peaks of gARID in gARID::GFP and gARID::GFP^C_*gsnf2*(−)_*pfg*(−)^ upstream of *calm* (top) and *rsph9* (bottom). Histograms show row read coverage of ChIP data at each base. Locations of TGTCT and TGCACA motifs are indicated as blue and green bars, respectively. The scales are shown in brackets. (B) Expression of CALM and RSPH9 in male and female gametocytes, observed using CALM::GFP and RSPH9::GFP parasites and fluorescence microscopy. Nuclei were stained with Hoechst 33342. (C) Fluorescence-activated cell sorting analysis of CALM::GFP (left) and RSPH9::GFP (right) before and after introducing mutations on the TGTCT and TGCACA motifs in their upstream peak regions. GFP-positive parasites were selected from cells gated with forward scatter and Hoechst. Nuclei were stained with Hoechst 33342.

The expression of the *gfp*-tagged genes was analyzed using FACS. Before introducing mutations, both CALM and RSPH9 were specifically expressed in the male gametocytes, which was consistent with the sex-specific transcriptome in which the genes are male-enriched (Fig 6B). For *calm*, the disruption of the TGTCT motif did not show a distinct change in the expression of CALM (Fig 6C, right). However, mutations in the TGCACA motifs resulted in a significant reduction (approximately two-fold) of the GFP signal intensity, and mutations in both motifs showed a similar result (Fig 6C, right). Therefore, this indicates that the TGCACA motif functions as a major *cis*-regulatory element for the activation of *calm*. For *rsph9*, mutations in the TGTCT motifs resulted in a significant reduction (approximately five-fold) of the GFP signal intensity (Fig 6 C, left). In contrast, mutations in the TGCACA motifs resulted in only a slight reduction of the signal (Fig 6 C, left), indicating a higher contribution of the TGTCT motifs on the activation of *rsph9* than that of the TGCACA motifs. Nevertheless, the parasites in which both motifs were mutated showed weaker signal intensity than did the TGTCT motif mutant (Fig 6C, left). These two reporter assays strongly suggested that the TGCACA motif functions as an independent *cis*-regulatory element for the activation of downstream genes. Furthermore, the effect of disrupting the TGTCT motifs and both of the two motifs in the reporter assays was similar to the level of downregulation in *gsnf2*(−) and *garid*(−), respectively. This result is consistent with the implication that the differences between the phenotypes of *garid*(−) and *gsnf2*(−) could be due to the function of gARID on the TGCACA motif.

## Discussion

In this study, we demonstrated the essential role of gARID in transcriptional regulation during the gametocyte development of *P. berghei*. The disruption of *garid* resulted in a complete loss of male gametocyte production and downregulation of most male-enriched genes, suggesting that gARID is a major regulator of male development. The ChIP-seq analyses showed that gARID is associated with gSNF2 at the TGTCT motif, which is a major *cis*-regulatory element for the activation of male genes. Along with the protein domain analysis of the ARID domain, this result suggested that gARID functions as a subunit of the SWI/SNF chromatin remodeling complex for regulating male genes. However, the disruption of the SWI/SNF complex on the TGTCT motif does not completely explain the phenotype of *garid*(−) in male development; *i.e.*, whereas *garid*(−) completely lost the ability to produce male gametocytes, *gsnf2*(−) still produced immature male gametocytes. In addition, differential expression analyses between *garid*(−) and *gsnf2*(−) showed a significant difference between their gametocyte transcriptomes. Through the ChIP-seq using gARID::GFP^C_*gsnf2*(−)_*pfg*(−)^, these differences were revealed to be derived from the function of gARID on the TGCACA motif. Therefore, we considered that gARID could be involved in two remodeling processes required for male gametocyte development, thereby associating with the two different chromatin remodeling complexes.

In female gametocytes, gARID is recruited to the female *cis*-regulatory element, TGTAYRTACA, which is associated with a female-specific transcription factor, PFG. However, in contrast to the marked phenotype in male gametocyte development, *garid*(−) barely showed abnormality in female gametocytes, which only resulted in decreased efficiency of ookinete maturation. This phenotype of *garid*(−) is very different from that of the *pfg*-knockout parasite, in which the development of fertilized parasites is arrested at the zygote stage (33). Therefore, we considered that gARID does not play an important role on the TGTAYRTACA motif, although recruited on it. In contrast, the target genes for the TGCACA motif (dKO targets), which are largely downregulated in *garid*(−), included a considerable number of female-enriched genes. Therefore, we speculate that the abnormality in female gametocytes of *garid*(−) is derived from the disruption of gARID function on the TGCACA motif.

The expression of gARID begins before the manifestation of each sex characteristic, which is earlier than that of the sex-specific transcriptional regulators, gSNF2 and PFG. This indicates that gARID functions before sex determination. Considering this and the roles of gARID on the TGCACA motif discussed above, we propose a model that describes the functions of gARID during gametocyte development (Fig 7). First, AP2-G triggers gametocytogenesis and activates gametocyte transcriptional regulator genes, including *garid*. Next, gARID activates early gametocyte genes on the TGCACA motifs, cooperating with a chromatin remodeler other than gSNF2. This step promotes early gametocyte development and preparation for male and female differentiation. In the male gametocyte lineage, male genes are activated via the SWI/SNF chromatin remodeling complex containing gSNF2 and gARID, associating with the male *cis*-regulatory element, TGTCT. Some of the genes activated by the TGCACA motif are further transcribed to be expressed male-specifically under the control of the TGTCT motif. In *garid*(−), the functions of these two *cis*-regulatory elements are both impaired, thus resulting in the complete absence of male gametocytes. In the female gametocyte lineage, transcriptional activators, AP2-FG and PFG, activate female genes, recruiting gARID. However, gARID probably plays no important role in activating these female genes. A subset of the TGCACA targets is also activated by AP2-FG and PFG for female-specific expression. In *garid*(−), these activators complement the function of TGCACA in part, and hence, female gametocyte development in *garid*(−) is only mildly affected.

**Fig 7.**
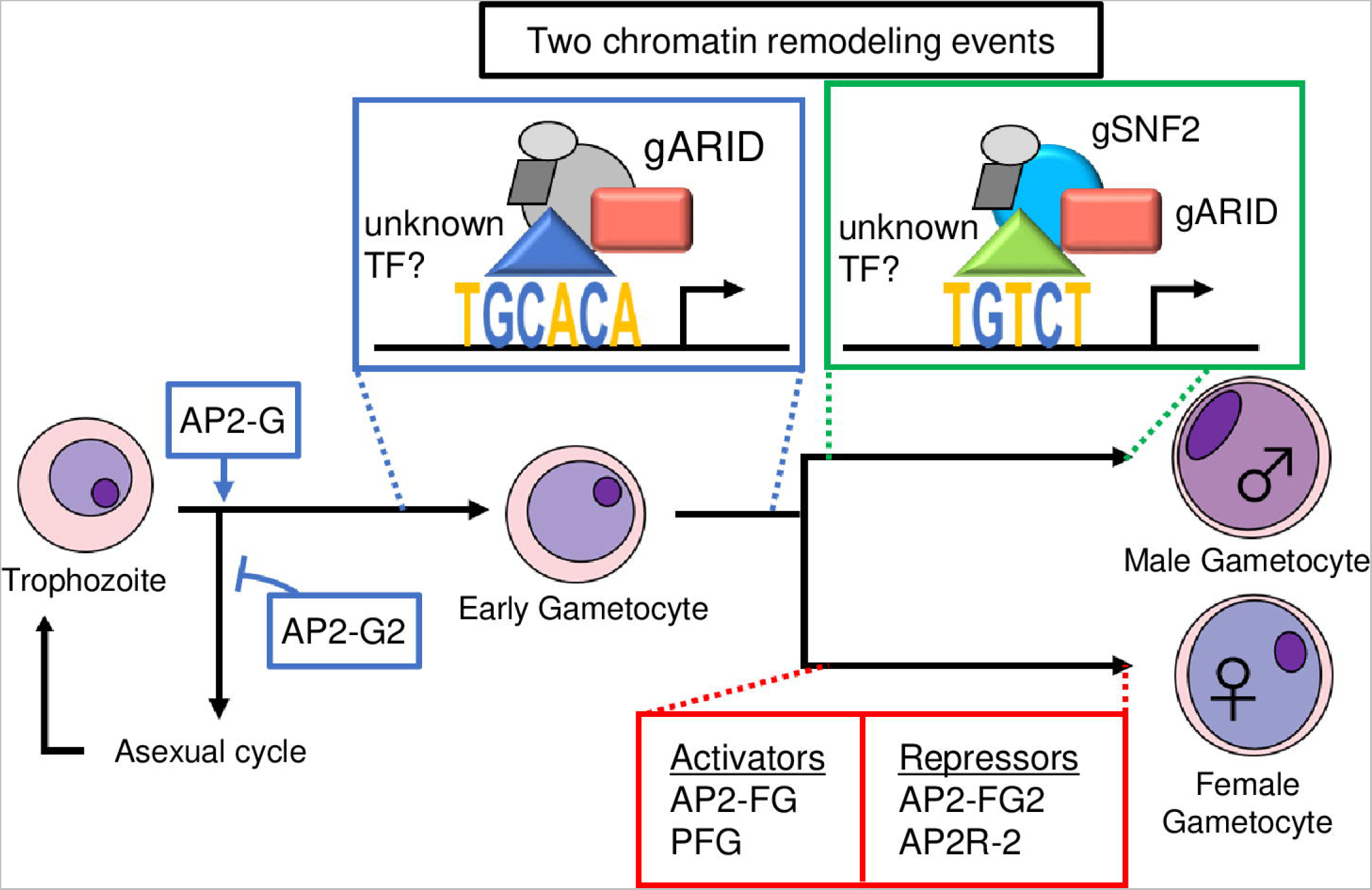
A schematic model for functions of gARID during gametocyte development in *Plasmodium berghei*. First, AP2-G triggers gametocytogenesis, followed by AP2-G2 supporting gametocyte differentiation through global gene repression. Next, gARID activates early gametocyte genes on the TGCACA motif, cooperating in a chromatin remodeling complex that does not contain gSNF2. After sex determination, male-gene transcription is further activated by the SWI/SNF chromatin remodeling complex containing gSNF2 and gARID, associating with the TGTCT motif. For female development, transcriptional activators (AP2-FG and PFG) and transcriptional repressors (AP2-FG2 and AP2R-2) promote female gametocyte maturation.

To conclude, we revealed the functions of gARID as a subunit of chromatin remodeling complexes. Moreover, our results showed that gARID functions with gSNF2 on the TGTCT and also with another, yet unknown, remodeler on the TGCACA motifs. Our results further showed that the chromatin remodeling events are important for both the male and female development of *P. berghei*. In a recent study, the disruption of *garid* ortholog in *P. falciparum* was also shown to affect both male and female development (44). Therefore, the gARID-mediated chromatin remodeling processes for early and male gametocyte development could be conserved in *Plasmodium* species. To deeply understand the chromatin remodeling process during the *Plasmodium* gametocyte development, further exploration of other transcriptional regulator genes is required. Most importantly, transcription factors that directly recognize the TGTCT and TGCACA motifs remain to be identified to date. Identifying these undiscovered factors can considerably help us understand the transcriptional regulation during gametocyte development, especially for early and male gametocyte development. As target genes of AP2-G contain most of the known sexual transcriptional regulator genes, we expect that these factors can be found in the AP2-G targets as well.

## Data Availability

All data produced in this study are fully available without restriction. All FASTQ files for ChIP-sequencing and RNA-sequencing experiments are available from the Gene Expression Omnibus database (accession numbers GSE235412).

## Acknowledgements

The authors are grateful to Asami Noro (Mie University) for technical assistance.

## Supplementary Data

**Fig S1. Alignment of amino acid sequences for gARID and its putative orthologs in Apicomplexan parasites by the ClustalW program in Mega X**

(*Plasmodium berghei*, gARID; *Plasmodium falciparum*, PF3D7_0603600; *Plasmodium knowlesi*, PKNH_1147100; *Cryptosporidium hominis*, XP_665701; *Eimeria tenella*, XP_013235762; *Toxoplasma gondii*, XP_018634911; *Neospora caninum*, XP_003885918; *Cystoisospora suis*, PHJ24466; and *Besnoitia besnoiti*, XP_029222326)

**Fig S2. Genotyping of transgenic parasites used for expression and phenotype analyses of *garid***

(A) gARID::mNG. (B) *garid*(−).

**Fig S3. Genotyping of transgenic parasites used for ChIP-seq experiments** (A) gARID::GFP. (B) gARID::GFP^C^. (C) gARID::GFP^C_*gsnf2*(−)^. (D) gARID::GFP^C_gsnf2(−)_pfg(−)^.

**Fig S4. Genotyping of transgenic parasites used for reporter assays**

(A) CALM::GFP. (B) RSPH9::GFP. (C) CALM::GFP^motif_mutated^. (D) RSPH9::GFP^motif_mutated^.

**Table S1. Differential expression analysis between *garid*(−) and wild type**

**Table S2. ChIP-seq analysis of gARID**

(A) Experiment 1. (B) Experiment 2. (C) Target genes.

**Table S3. Differential expression analysis between *garid*(−) and *gsnf2*(−)**

**Table S4. ChIP-seq analysis of gARID using gARID::GFP^C_*gsnf2*(−)_*pfg*(−)^**

(A) Experiment 1. (B) Experiment 2. (C) dKO targets.

**Table S5. List of primers used in this study**

