## Supplementary figures and images for "gARID-associated chromatin remodeling events are essential for gametocyte development in *Plasmodium*"

### Figure S1

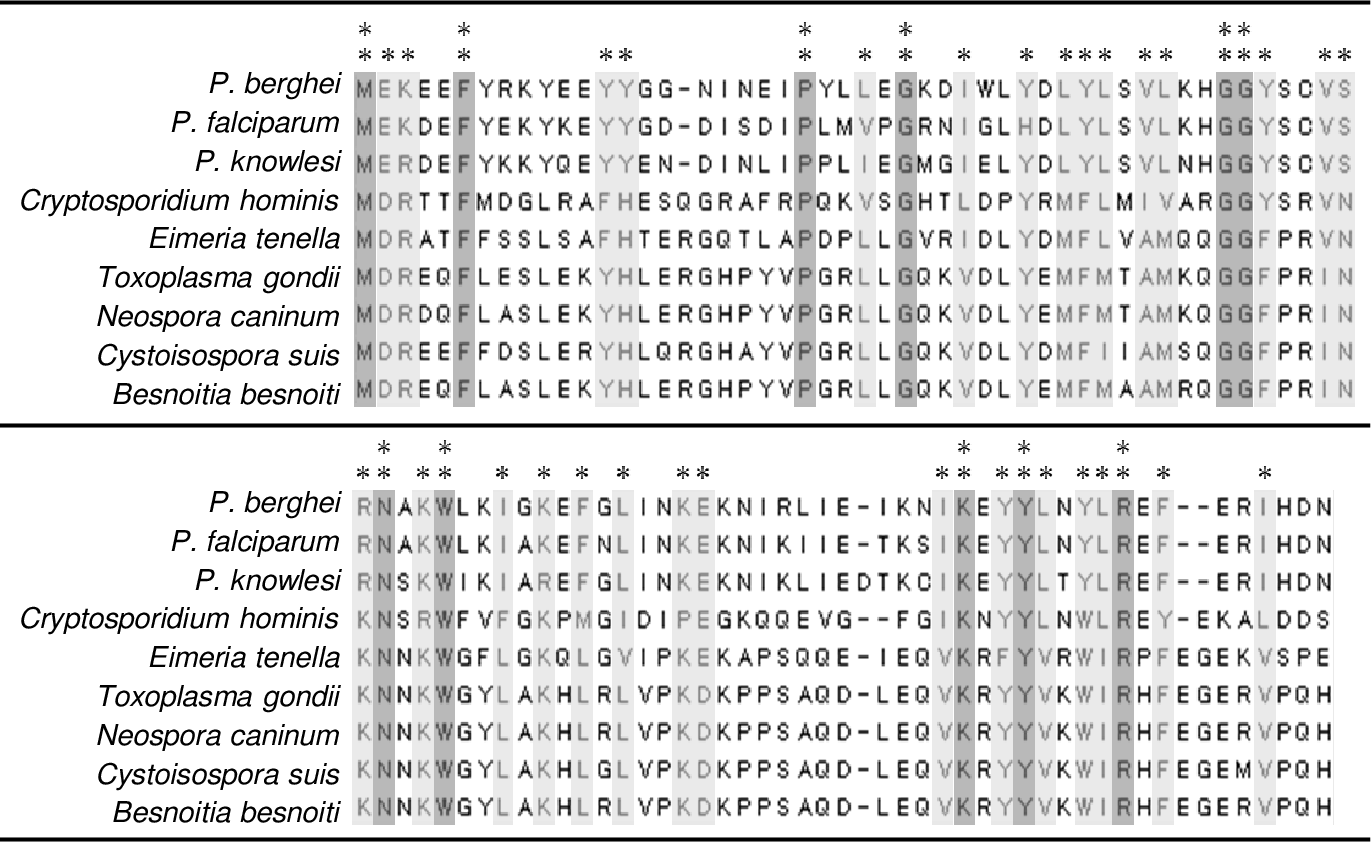

### Figure S2

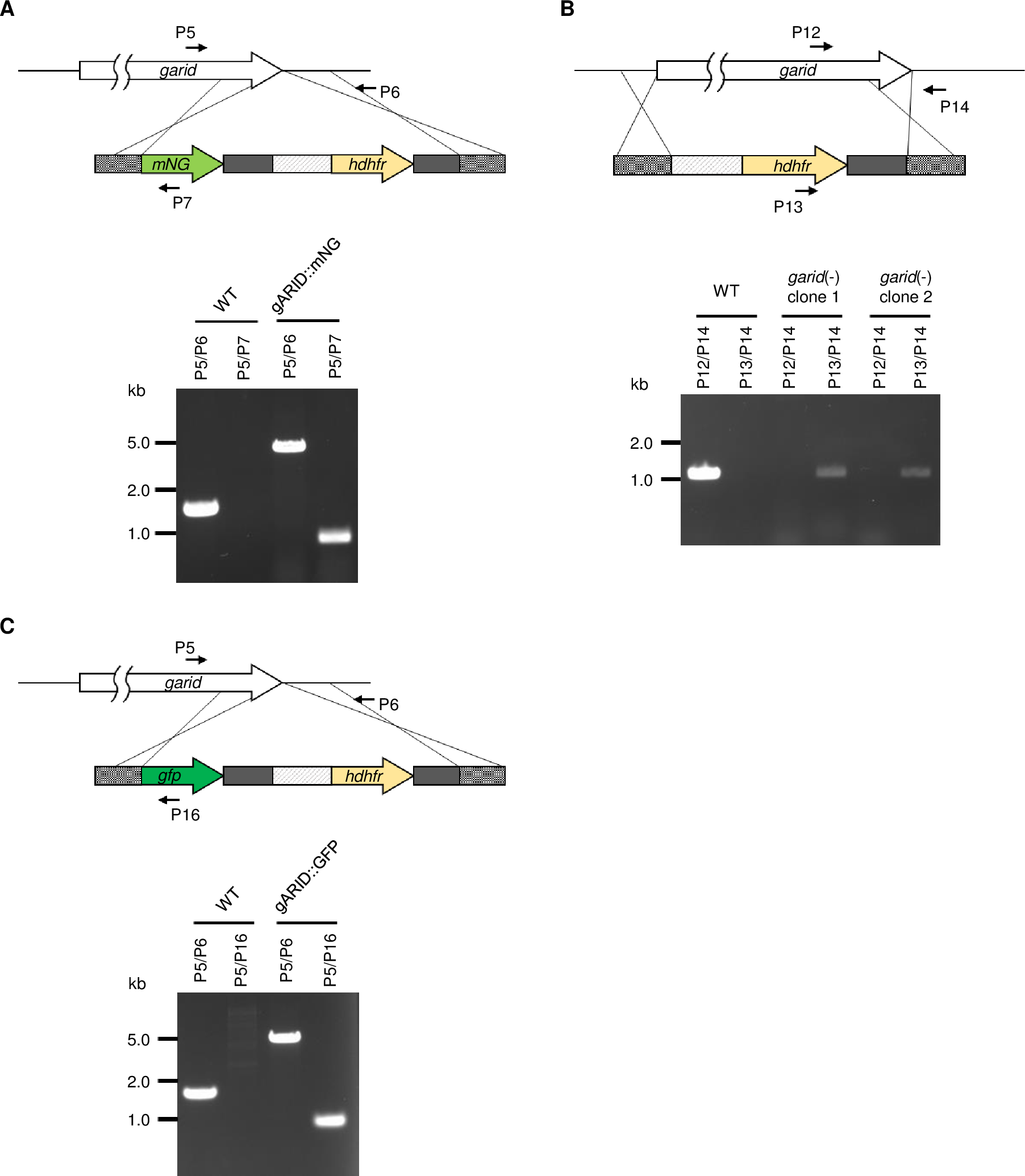

### Figure S3

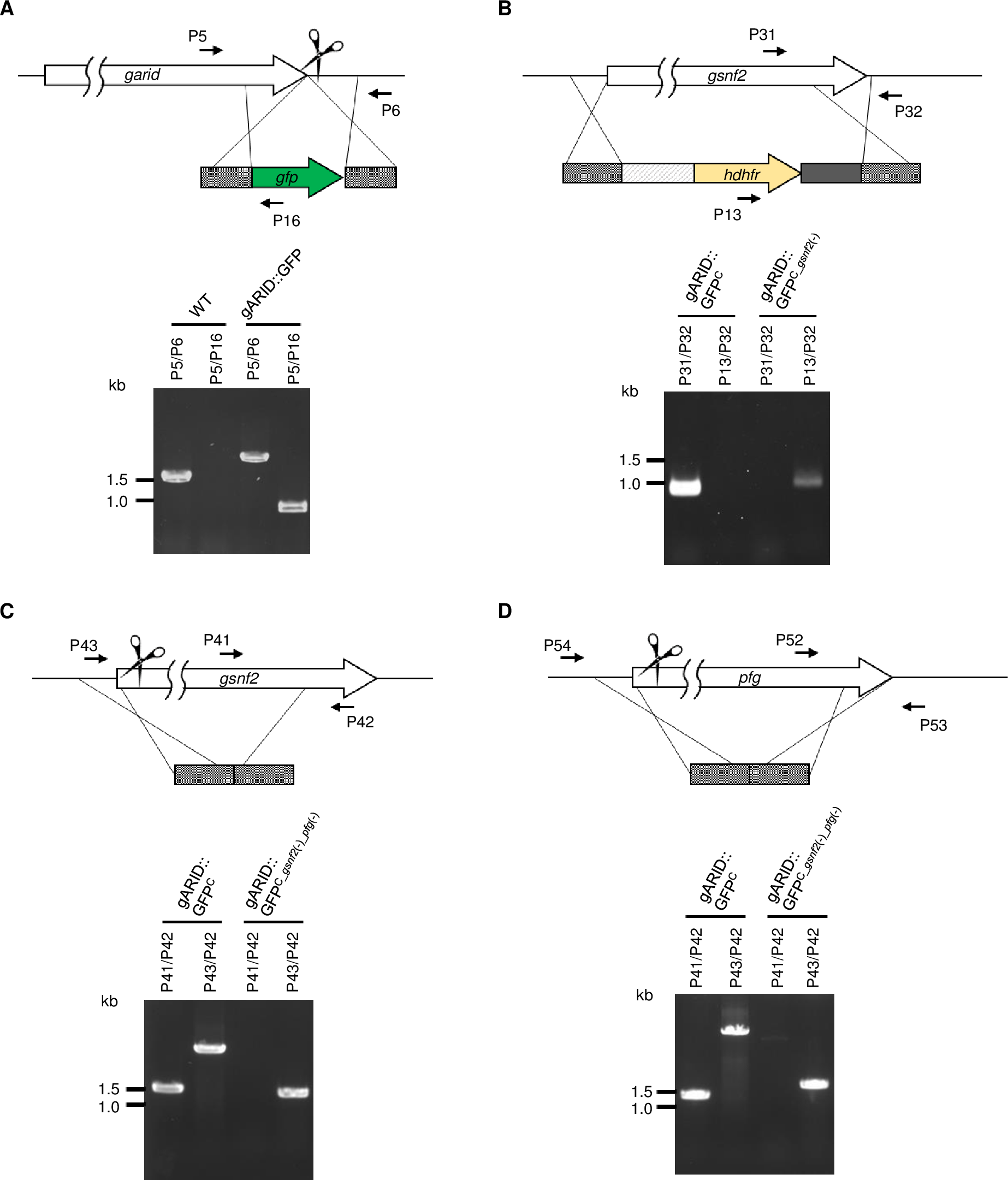

### Figure S4

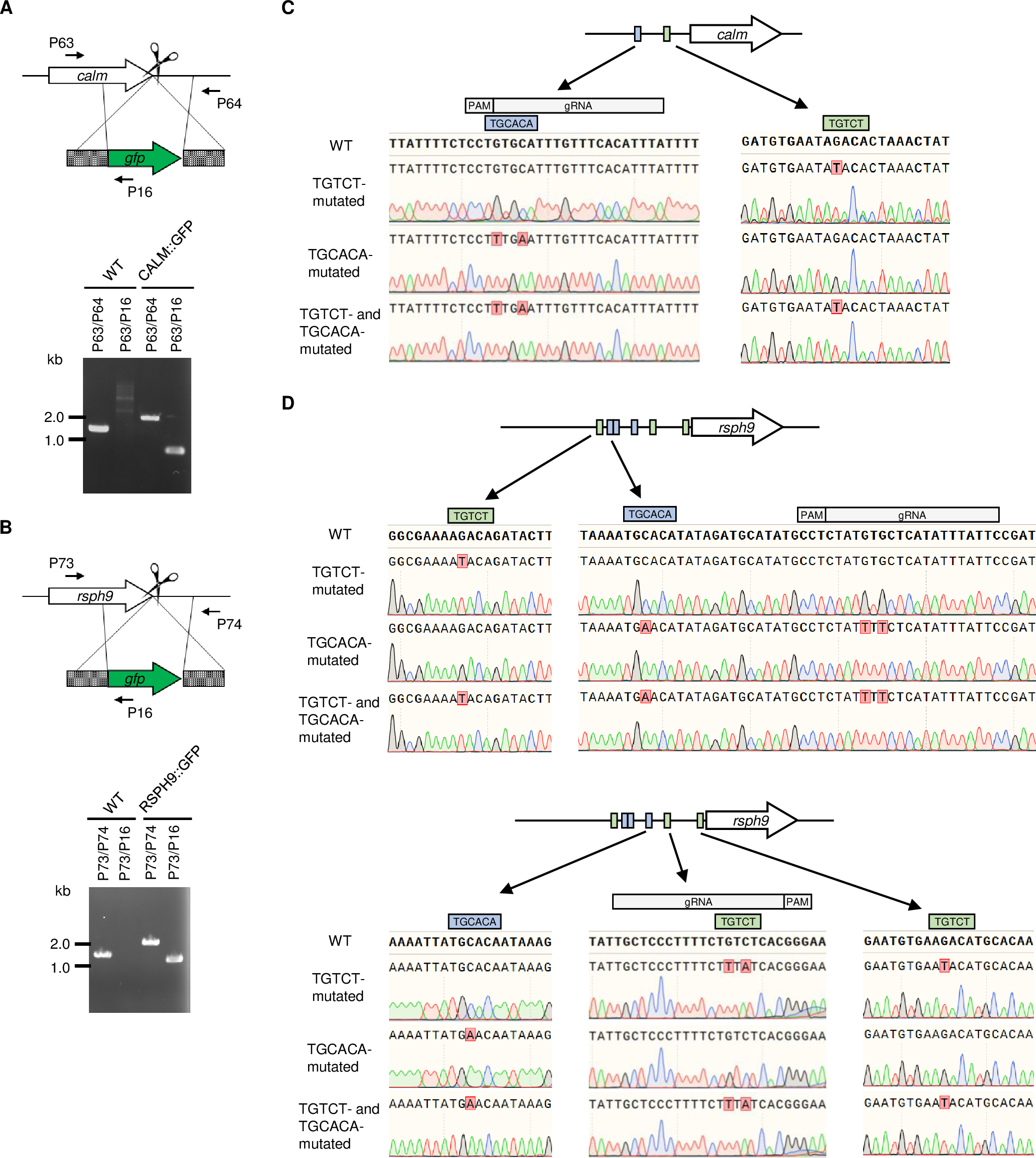
